# RabA- and RabE-mediated trafficking define distinct branches of plant pre-invasive immunity

**DOI:** 10.64898/2026.11.10.765595

**Authors:** Mads Eggert Nielsen, Jelco Mensch, David Havndrup Munch, Héctor Molinelli Rubiato

**Author notes:** Corresponding author: Mads Eggert Nielsen.

## Abstract

Pre-invasive immunity constitutes a major barrier to disease and protects plants against the vast majority of filamentous fungal and oomycete pathogens through localized defence responses at sites of attempted invasion. Although membrane trafficking is essential for these responses, demonstrated by the requirement for the syntaxins PEN1 and SYP122, the underlying trafficking pathways remain poorly understood. Here, we show that trafficking mediated by the RabE and RabA GTPase subfamilies defines two functionally distinct branches of plant pre-invasive immunity. Inhibition of RabE rapidly compromised immunity to powdery mildew fungi, including the durable immunity conferred by *mlo* mutations, and reduced focal accumulation of PEN1 at papillae without substantially affecting callose deposition. By contrast, inhibition of RabA disrupted plasma membrane localization of multiple proteins and preferentially compromised immunity to the oomycete pathogen *Phytophthora infestans*. Together, these findings support a model in which RabE promotes the powdery mildew-specific function of PEN1 in pre-invasive immunity, whereas RabA contributes to a broader defence pathway associated with the shared function of PEN1 and SYP122. Our results support a model in which pre-invasive immunity is coordinated by specialized trafficking modules with distinct pathogen requirements and establish RabA-and RabE-mediated trafficking as defining components of these pathways.

## Introduction

Filamentous fungal and oomycete pathogens invade plant tissues by breaching the epidermal cell wall and are responsible for some of the most destructive diseases in natural and agricultural ecosystems (Savary et al., 2019). To counter invasion, plants have evolved highly effective pre-invasive defence mechanisms that are activated upon pathogen attack (Nielsen, 2024). These responses are characterized by the focal assembly of papillae beneath attempted penetration sites and the formation of encasements around invasive structures following successful ingress (O’Connell and Panstruga, 2006). Together, these cell-autonomous defence structures contribute to the remarkable durability of nonhost resistance towards a wide range of filamentous pathogens (Panstruga and Moscou, 2020). The importance of these responses is underscored by the observation that loss of the two closely related SYP12-clade syntaxins PENETRATION1 (PEN1; syntaxin in plants121, SYP121) and SYP122 abolishes papilla and encasement formation and compromises immunity to phylogenetically diverse fungal and oomycete pathogens in *Arabidopsis thaliana* (Rubiato et al., 2022).

PEN1 and SYP122 are members of the soluble N-ethylmaleimide-sensitive factor attachment protein receptor (SNARE) family, and their requirement for papilla and encasement formation demonstrates that these defence structures depend on polarized membrane trafficking. Although PEN1 and SYP122 share an overlapping role in the assembly of these defence structures and in the establishment of broad-spectrum immunity, genetic analyses have revealed an additional specialized function of PEN1 during interactions with powdery mildew fungi (Rubiato et al., 2022). In contrast to SYP122, PEN1 is required for rapid pre-invasive immunity towards powdery mildew fungi and has been proposed to participate in a highly dynamic trafficking route that enables focal delivery of defence cargo to sites of fungal attack (Collins et al., 2003; Assaad et al., 2004). This functional distinction suggests that general pre-invasive immunity and powdery mildew-specific immunity may rely on partially separate membrane trafficking pathways delivering the anti-microbial cargo that blocks the pathogen (Nielsen and Thordal-Christensen, 2013; Hansen and Nielsen, 2017; Nielsen, 2024).

Consistent with this idea, we recently proposed that pre-invasive immunity comprises at least two genetically separable trafficking pathways defined by specific ADP-ribosylation factor guanine nucleotide exchange factors (ARF-GEFs) required for vesicle budding (Richter et al., 2014; Brumm et al., 2020). A broadly acting pathway, associated with the shared immune function of PEN1 and SYP122, requires BIG1-BIG4 and contributes to immunity against diverse filamentous pathogens (Munch *et al*., 2026). In contrast, a GNOM-dependent pathway is specifically associated with the specialized function of PEN1 during powdery mildew attack (Nielsen et al., 2012; Nielsen et al., 2017). These findings raise the possibility that distinct membrane trafficking regulators define general and powdery mildew-specific immune pathways. However, the molecular machinery acting downstream of BIG1-BIG4 and GNOM remains largely unknown.

Rab GTPases are master regulators of vesicle trafficking and membrane identity that help ensure correct SNARE assembly and thereby confer specificity to cargo delivery (Stenmark, 2009). In plants, extensive genetic redundancy within Rab subfamilies has often hindered the identification of individual members that contribute to specific membrane trafficking pathways (Nielsen, 2020). Instead, dominant-negative Rab variants have provided important insights into subfamily functions. In this way, members of the RabA and RabE subfamilies have been shown to regulate secretion, recycling, and polarized membrane transport at the plasma membrane, processes that are central to the delivery of defence cargo during pathogen attack (Zheng et al., 2005; Chow et al., 2008; Speth et al., 2009; Orr et al., 2021; Pang et al., 2022; Koch et al., 2026). In support of this, both RabA and RabE dependent secretion is required for resistance against different species of the pathogenic oomycete *Phytophthora* (Tomczynska et al., 2018; Li et al., 2022; Pandey et al., 2021; Yuen et al., 2024). However, whether RabA and RabE GTPases directly control pre-invasive immunity, and how they connect to the shared and specialized immune functions of PEN1 and SYP122, remains unknown.

Here, we show that RabA- and RabE-mediated trafficking define distinct branches of plant pre- invasive immunity. RabE-dependent trafficking is specifically required for immunity to powdery mildew fungi and contributes to durable MILDEW RESISTANCE LOCUS O (*mlo*)-mediated immunity, whereas RabA-dependent trafficking preferentially contributes to immunity against the oomycete pathogen *Phytophthora infestans*. These findings identify RabA and RabE as defining components of two genetically and functionally distinct immune trafficking pathways. We propose that RabE acts in a pathway associated with the specialized function of PEN1 during powdery mildew attack, whereas RabA contributes to a broader immune pathway linked to the shared function of PEN1 and SYP122. Together, our results provide a mechanistic framework for understanding how plants deploy specialized membrane trafficking routes to counter diverse filamentous pathogens.

## Results

### RabA-mediated trafficking contributes to pre-invasive immunity only after prolonged inhibition

Recently, RabA2a was directly linked to PEN1 and SYP122 and found required for their proper localization suggesting a role in defence-associated membrane trafficking (Pang et al., 2022). To assess a potential role for RabA in pre-invasive immunity against non-adapted powdery mildew fungi, we employed dexamethasone (DEX)-inducible dominant-negative variants of RabA2a previously described by Chow et al. (2008). Constitutively active RabA2a (QL) plants displayed severe developmental defects even in the absence of induction and were therefore excluded from subsequent analyses. We initially induced expression of the dominant-negative alleles RabA2a (SN) and RabA2a (NI) for 3 h before inoculation with *Blumeria hordei*. Under these conditions, penetration frequencies did not differ significantly from those of control plants (Supplementary Fig. 1a). By contrast, repeated DEX treatments administered 48 h, 24 h and 3 h before inoculation resulted in a marked increase in fungal penetration, demonstrating that prolonged inhibition of RabA compromises pre-invasive immunity (Fig. 1a). RabA2a (NI) caused a stronger phenotype than RabA2a (SN), consistent with previous reports (Chow et al., 2008). Notably, prolonged induction of RabA2a (NI) also impaired callose accumulation at both unsuccessful penetration attempts (papillae) and successful penetration sites (encasements) (Fig. 1b-g; Supplementary Fig. 1b). The resulting phenotype closely resembled the defective cell wall responses previously described in *pen1 syp122* mutants (Rubiato et al., 2022). Together, these data indicate that RabA function contributes to powdery mildew immunity but that substantial inhibition is required before defects in pre-invasive immunity and callose deposition become apparent.

**Fig. 1:**
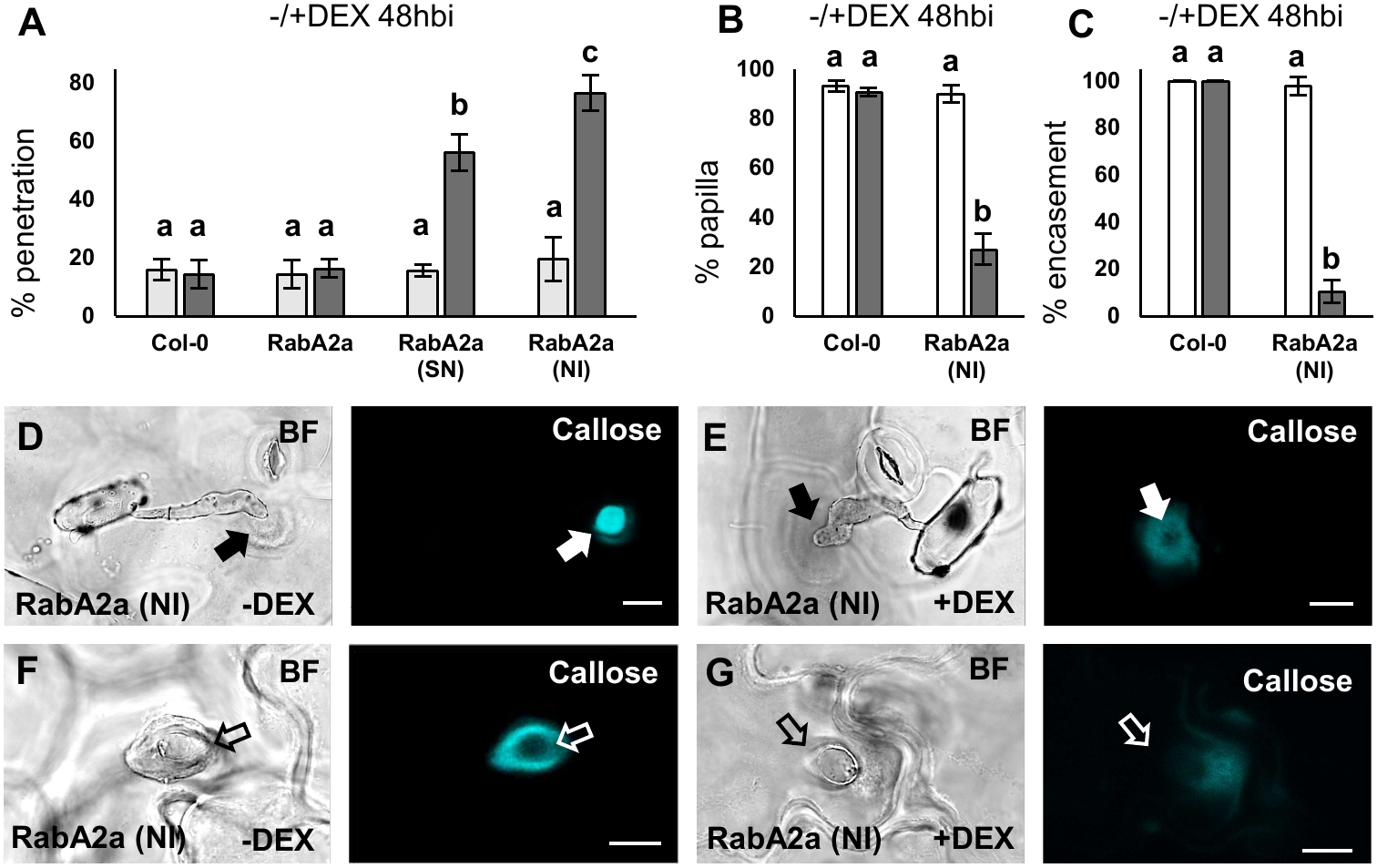
Delayed effects of RabA inhibition on powdery mildew pre-invasive immunity. Frequency of successful penetrations of leaf epidermal cells by *B. hordei* in Col-0, *DEX::RabA2a*, *DEX::RabA2a (SN)* or *DEX::RabA2a (NI)* sprayed with either control (white bars) or 20 µM dexamethasone (grey bars) 48, 24 and 3 hours before inoculation (A). Frequency of papillae (B) and encasements (C) in response to *B. hordei* as detected by staining for callose in Col-0 or *DEX::RabA2a (NI)* treated as in A. Confocal images of callose accumulations in response to *B. hordei* attack (arrows) in non-penetrated (D,E) and penetrated (F,G) cells (open arrows point to the forming haustorium) in *DEX::RabA2a (NI)* treated with either control (D,F) or 20 µM dexamethasone (E,G). Scale bars = 10 µm. All values are mean ±SD. *P* < 0.05, one-way ANOVA and Tukey HSD test, n = 4-5.

### RabA maintains plasma membrane localization of selected proteins

The requirement for prolonged RabA inhibition to affect pre-invasive immunity contrasts with reports that RabA disruption rapidly alters PEN1 localization (Pang et al., 2022). To investigate this discrepancy, we monitored GFP-PEN1 after induction of RabA2a (NI). Strikingly, GFP-PEN1 localization was already strongly affected 24 h after induction, well before measurable defects in pre-invasive immunity were observed (Fig. 2a). Following prolonged induction, PEN1 became progressively depleted from the plasma membrane and instead accumulated on intracellular membranes frequently associated with fragmented vacuoles (Fig. 2b). Co-expression with the tonoplast marker Vesicle-associated membrane protein 711 (VAMP711) confirmed substantial relocalization of PEN1 to vacuolar membranes, whereas no comparable accumulation was detected in compartments marked by ARA7 or 2xFYVE (Supplementary Fig. 2).

**Fig. 2:**
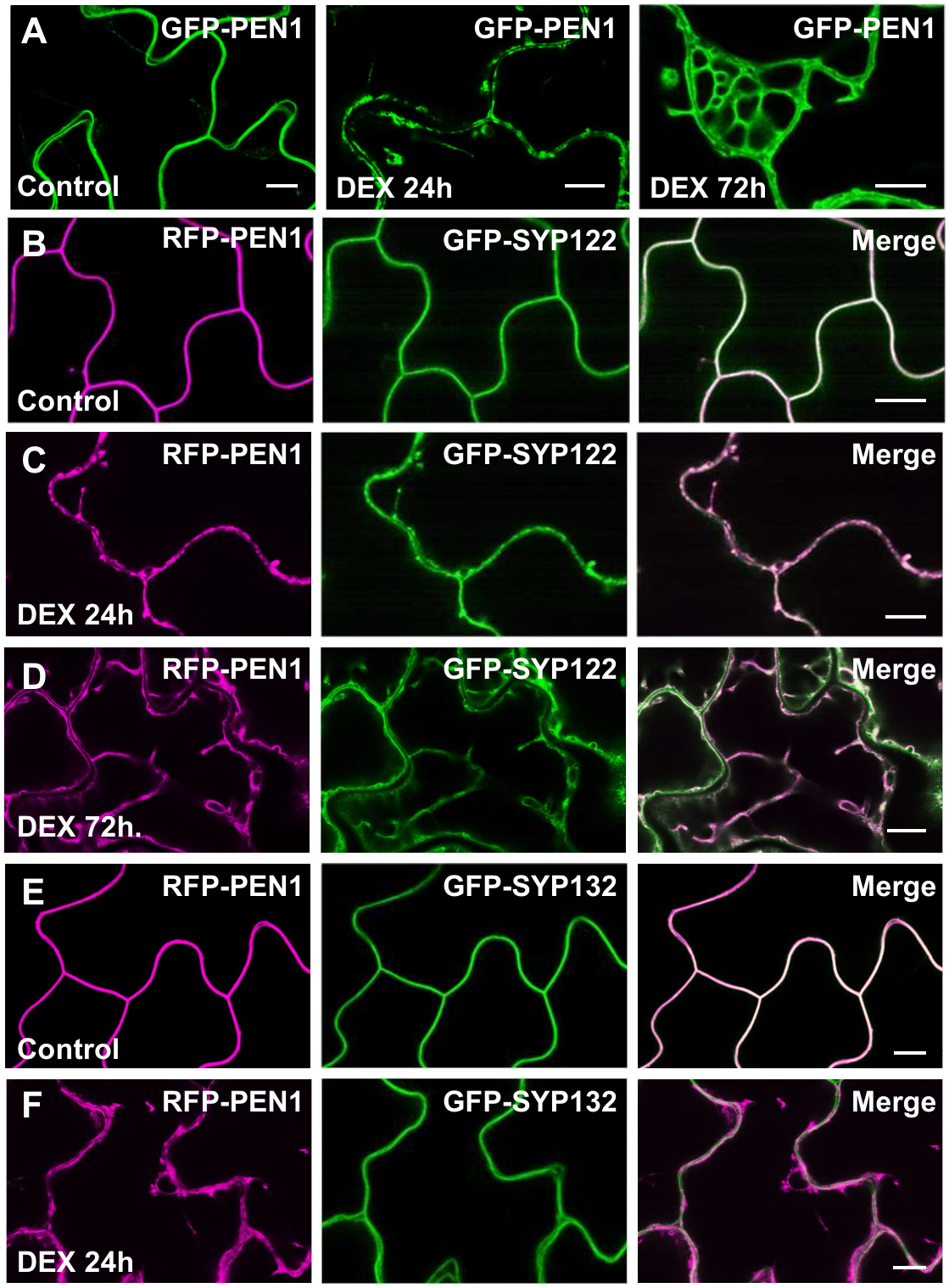
RabA inhibition depletes PEN1 and SYP122 from the plasma membrane. Representative confocal micrographs of epidermal leaf cells from 7-day-old *DEX::RabA2a (NI)* seedlings grown on medium containing 20 μM dexamethasone for the indicated periods before imaging. GFP-PEN1 localization following induction of RabA2a (NI) (A). Co-localization of RFP-PEN1 and GFP-SYP122 (B-D). Co-localization of RFP-PEN1 and GFP-SYP132 (E,F). Expression of RabA2a (NI) progressively altered the localization of PEN1 and SYP122, whereas SYP132 localization remained largely unaffected. Scale bars, 10 μm.

A similar temporal pattern was observed for GFP-SYP122 (Fig. 2b-d). Moreover, RabA2a (NI) altered the localization of several additional plasma membrane-associated proteins, including MLO2, Tetraspanin8 (TET8) and PEN3, within 24 h of induction (Supplementary Fig. 3a-f). In contrast, the plasma membrane syntaxin SYP132 remained largely unaffected, and secretion of secRFP was only weakly perturbed even after prolonged induction (Fig. 2e,f; Supplementary Fig. 3g,h).

These observations demonstrate that inhibition of RabA rapidly disrupts the plasma membrane localization of a specific subset of proteins, including several previously implicated in plant immunity. Yet these trafficking defects preceded detectable reductions in pre-invasive immunity, revealing a striking temporal disconnect between plasma membrane protein localization and penetration resistance. This finding suggests that RabA primarily maintains plasma membrane homeostasis rather than directly mediating immune trafficking. We therefore asked whether other Rab GTPases contribute more directly to pre-invasive immunity.

### RabE is required for immunity against non-adapted powdery mildew fungi

RabE proteins are plant homologues of the Rab8 family that promote secretion toward the plasma membrane (Zheng et al., 2005; Koch et al., 2026). To examine whether RabE-dependent trafficking contributes to pre-invasive immunity, we analysed YFP-RabE1d, which localized predominantly to intracellular puncta distinct from mCherry-RabA2a-positive structures with some overlap at the plasma membrane (Supplementary Fig. 4a,b). In response to attack by *B. hordei*, RabA2a and RabE1d showed only little enrichment in papillae or encasements, suggesting functions upstream of focal cargo delivery rather than accumulating directly within defence structures (Fig. 3a,b).

**Fig. 3:**
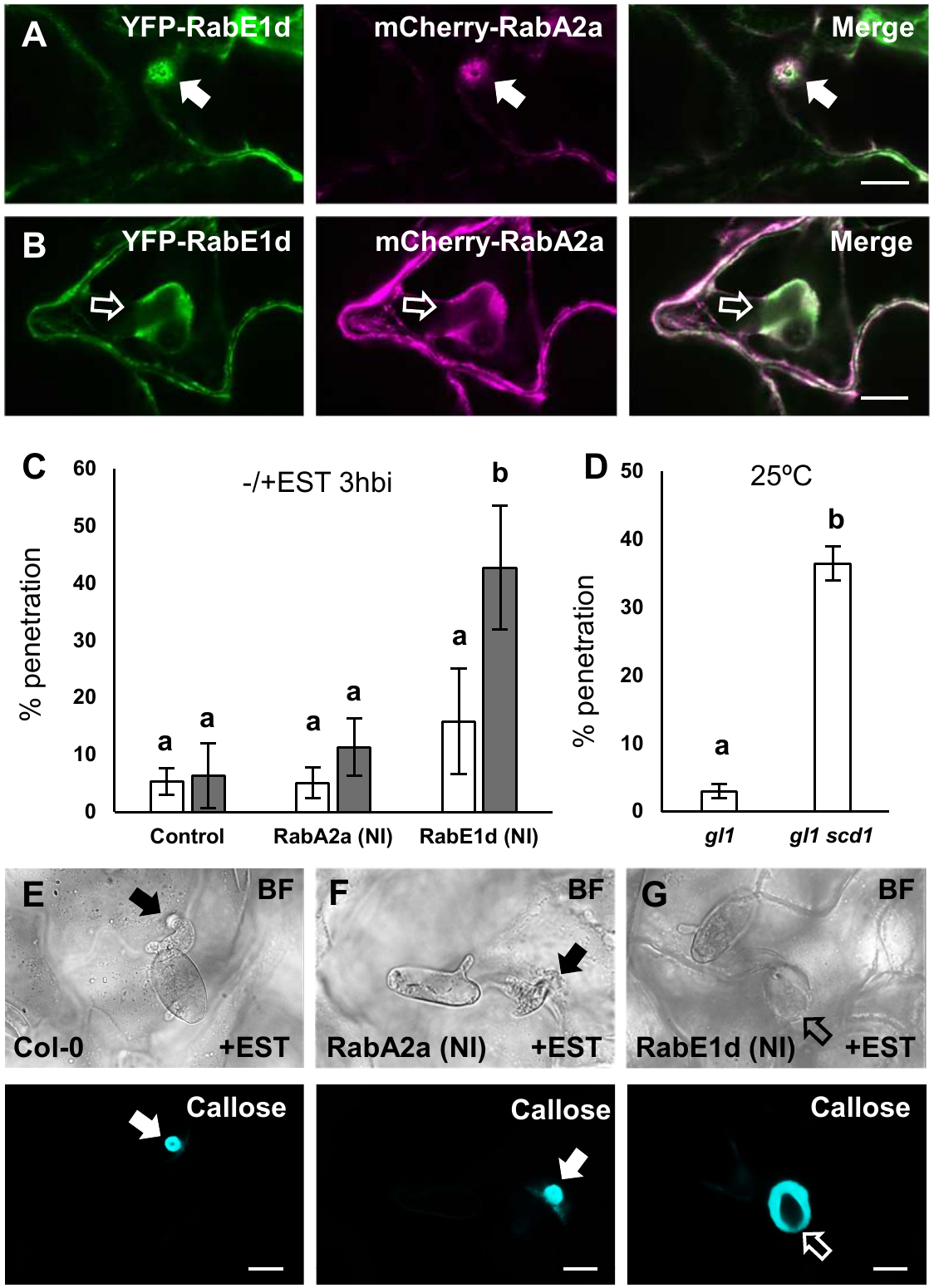
Rapid impairment of pre-invasive immunity following RabE inhibition. Localization of YFP-RabE1d and mCherry-RabA2a in response to *B. hordei* attack (arrows) in non-penetrated (A) and penetrated (B) cells (open arrows point to the forming haustorium). Frequency of successful penetrations of leaf epidermal cells by *B. hordei* in Col-0, *EST::RabA2a (NI)* or *EST::RabE1d (NI)* sprayed with either control (white bars) or 20 µM *β*-estradiol (grey bars) 3 hours before inoculation (C). Frequency of successful penetrations of leaf epidermal cells by *B. hordei* in *glabrous1* (*gl1*) or *gl1 scd1* at 25°C (D). Accumulation of callose in response to *B. hordei* attack (arrows) in non-penetrated (E,F) and penetrated (G) cells (open arrows point to the forming haustorium) in Col-0, *EST::RabA2a (NI)* or *EST::RabE1d (NI)* sprayed with 20 µM *β*-estradiol 3 hours before inoculation. Scale bars, 10 μm. All values are mean ±SD. *P* < 0.05, one-way ANOVA and Tukey HSD test, n = 4-5.

To compare the roles of RabA and RabE directly, we generated estradiol (EST)-inducible dominant-negative RabE1d (NI) and RabA2a (NI) lines in a GFP-PEN1 background. In line with previous reports, prolonged induction of either construct severely affected plant growth (Supplementary Fig. 4c). However, unlike RabA2a (NI), induction of RabE1d (NI) for 24 h did not visibly alter GFP-PEN1 localization (Supplementary Fig. 4d).

Despite the absence of obvious PEN1 trafficking defects, induction of RabE1d (NI) only 3 h before inoculation significantly increased penetration frequencies of *B. hordei*, whereas equivalent induction of RabA2a (NI) had no detectable effect (Fig. 3c). Consistent with this finding, the temperature-sensitive mutation stomatal cytokinesis-defective 1-1 (*scd1*), affecting a RabE guanine nucleotide exchange factor, likewise compromised pre-invasive immunity at the restrictive temperature (Fig. 3d). Unexpectedly, neither RabE1d (NI) expression nor loss of SCD1 substantially affected pathogen-induced callose deposition (Fig. 3e-g; Supplementary Fig. 5). Thus, RabE-dependent trafficking contributes directly to pre-invasive immunity against non-adapted powdery mildew fungi and can be genetically uncoupled from callose deposition.

### RabE promotes focal accumulation of PEN1 at sites of fungal attack

Because callose deposition remained largely intact following RabE inhibition, we investigated the recruitment of defence-associated plasma membrane proteins to attack sites. In wild-type plants, both PEN1 and PEN3 accumulated at papillae, although PEN1 exhibited a substantially higher papilla-to-plasma membrane fluorescence ratio (Fig. 4). Expression of RabA2a (NI) reduced PEN1 accumulation at papillae, but interpretation was complicated by broader defects in plasma membrane localization. In contrast, dominant-negative RabE1d produced a more selective phenotype. PEN1 accumulation at papillae was consistently reduced, whereas PEN3 recruitment was comparatively unaffected (Fig. 4). Similar defects were observed in *scd1* mutants (Supplementary Fig. 6a-c). Notably, PEN1 accumulation at encasements remained largely unaffected, indicating that distinct trafficking pathways contribute to PEN1 recruitment during different stages of defence (Supplementary Fig.6d-f). Together, these data identify PEN1 focal accumulation, rather than callose deposition, as the cellular process most closely associated with RabE-dependent pre-invasive immunity.

**Fig. 4:**
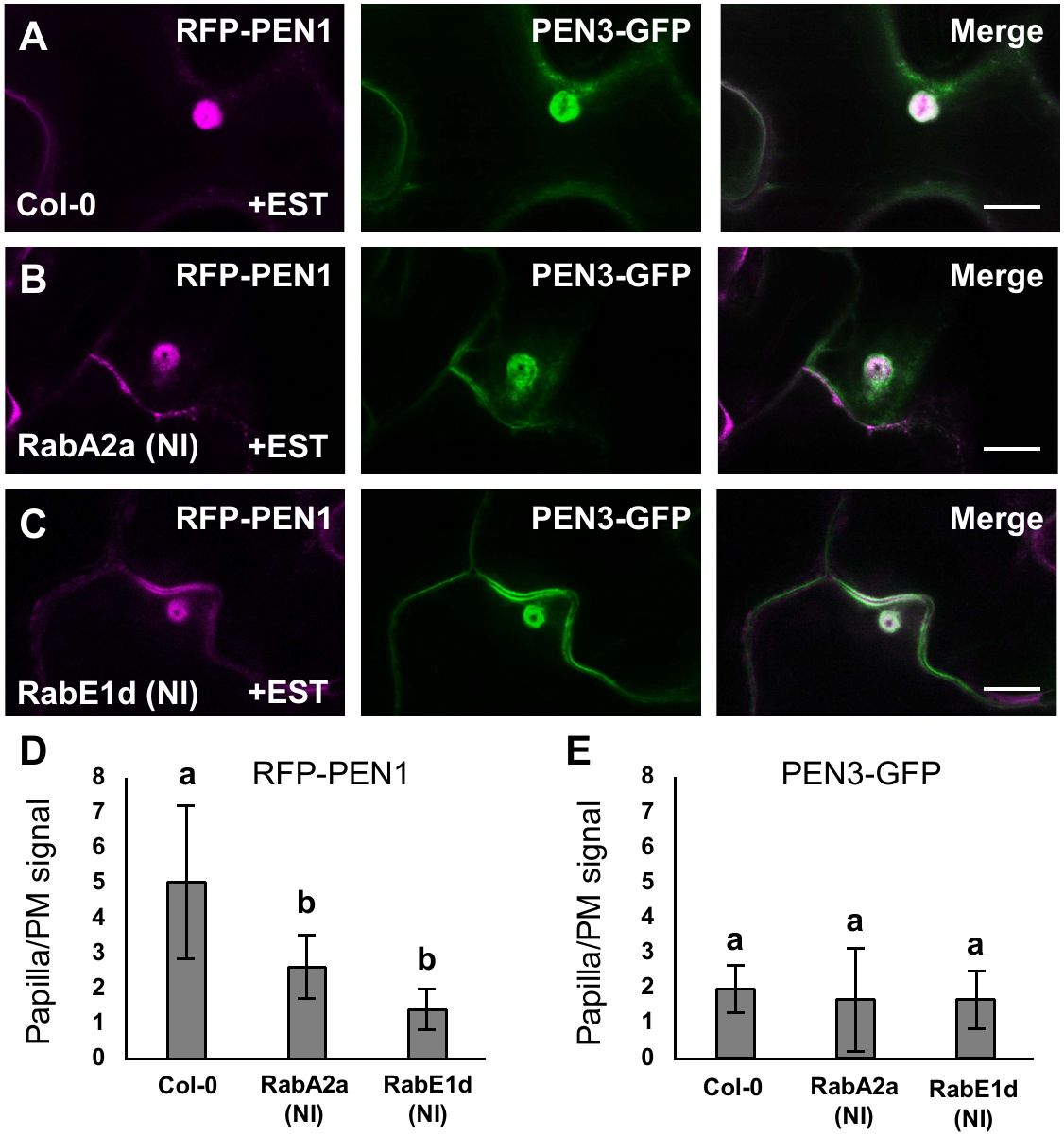
RabE preferentially affects PEN1 accumulation at papillae. Papillary accumulation of RFP-PEN1 and GFP-PEN3 in response to *B. hordei* attack in non-penetrated and penetrated cells in Col-0 (A), *EST::RabA2a (NI)* (B) or *EST::RabE1d (NI)* (C) sprayed with 20 µM *β*-estradiol 3 hours before inoculation. Quantification of relative papilla to plasma membrane (PM) signal for RFP-PEN1 (D) and PEN3-GFP (E). Scale bars, 10 μm. All values are mean ±SD. *P* < 0.05, one-way ANOVA and Tukey HSD test, n = 15-18.

### RabA and RabE regulate distinct branches of pre-invasive immunity

Previously, we proposed that plant pre-invasive immunity towards filamentous pathogens involves at least two mechanistically distinct trafficking pathways, where one acts broadly against fungi and oomycetes, and the other is dedicated towards powdery mildew fungi specifically (Rubiato et al., 2022; Munch et al., 2026). To determine whether RabA and RabE contribute differentially to these pre-invasive immunity pathways, we challenged induced plants with spores from the oomycete pathogen *P. infestans*. In contrast to the strong effects observed during interaction with *B. hordei*, inhibition of RabE did not increase penetration by *P. infestans*. In contrast, RabA2a(NI) markedly compromised immunity to the oomycete pathogen, demonstrating that RabA- and RabE-mediated trafficking underpin distinct branches of pre-invasive immunity (Fig. 5a, Supplementary Video 1). Likewise, shifting *scd1* plants to the restrictive temperature did not alter penetration rates of *P. infestans* (Fig. 5b). Despite enhanced pathogen entry, host cells expressing RabA2a (NI) retained the ability to mount localized defence responses. (Fig. 5c-e). These findings establish that RabA- and RabE-mediated trafficking pathways make differential contributions to pre-invasive immunity against non-adapted fungal and oomycete pathogens.

**Fig. 5:**
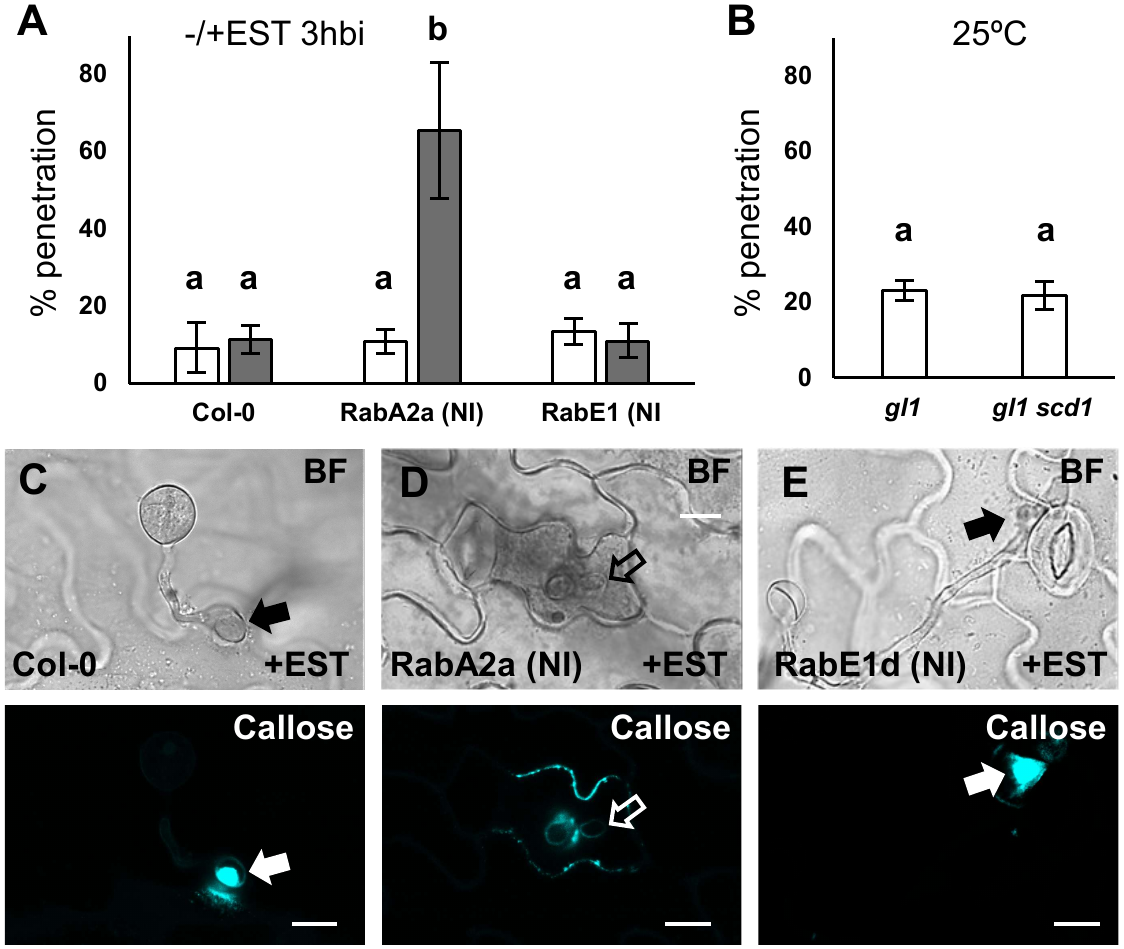
Differential requirements for RabA and RabE during oomycete invasion. Frequency of successful penetrations of leaf epidermal cells by *P. infestans* in Col-0, *EST::RabA2a (NI)* or *EST::RabE1d (NI)* sprayed with either control (white bars) or 20 µM *β*-estradiol (grey bars) 3 hours before inoculation (A). Frequency of successful penetrations of leaf epidermal cells by *P. infestans* in *glabrous1* (*gl1*) or *gl1 scd1* Scale bars, 10 μm (B). Accumulation of callose in response to *P. infestans* attack (arrows) in non-penetrated (C,F) and penetrated (D) cells (open arrows point to the forming haustorium) in Col-0, *EST::RabA2a (NI)* or *EST::RabE1d (NI)* sprayed with 20 µM *β*-estradiol 3 hours before inoculation. Scale bars, 10 μm. All values are mean ±SD. *P* < 0.05, one-way ANOVA and Tukey HSD test, n = 4-5.

### RabE-dependent trafficking contributes to *mlo*-based powdery mildew immunity

Recently, we have shown that the near complete immunity to the adapted powdery mildew fungus *Golovinomyces orontii* in the *mlo2 mlo6 mlo12* mutant requires PEN1 and SYP122 (Munch et al., 2026). To assess whether Rab-dependent trafficking also contributes to this durable *mlo*-mediated immunity, we introduced inducible RabA2a (NI) and RabE1d (NI) into the highly resistant *mlo2 mlo6 mlo12* background. Whereas inhibition of RabA had no measurable impact on pre-invasive immunity to the adapted powdery mildew fungus *G. orontii*, expression of RabE1d (NI) significantly increased successful penetration events (Fig. 6a).

**Fig. 6:**
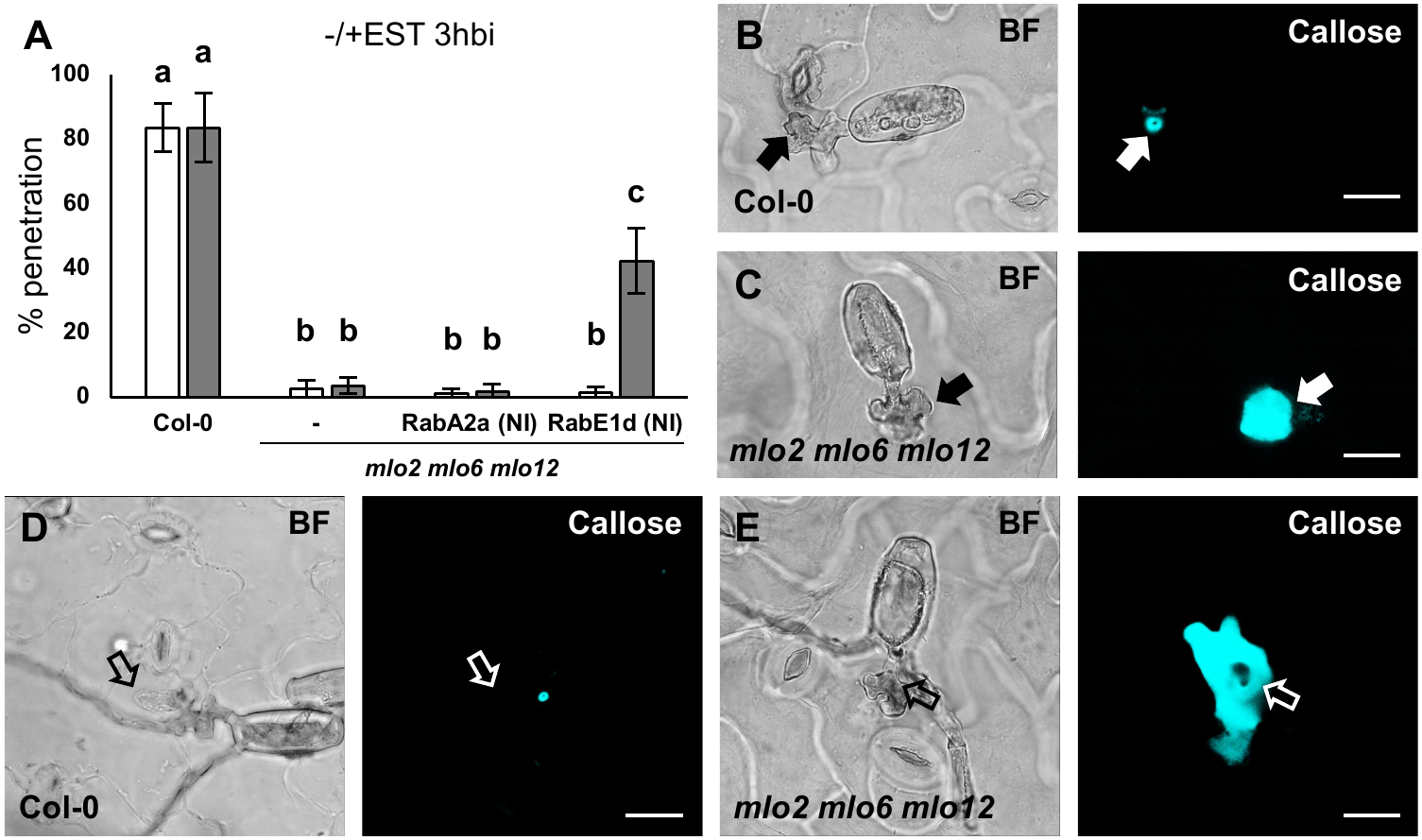
RabE inhibition partially overcomes *mlo*-based immunity. Frequency of successful penetrations of leaf epidermal cells by *G. orontii* in Col-0, *mlo2 mlo6 mlo12, mlo2 mlo6 mlo12 EST::RabA2a (NI)* or *mlo2 mlo6 mlo12 EST::RabE1d (NI)* sprayed with either control (white bars) or 20 µM *β*-estradiol (grey bars) 3 hours before inoculation (A). Accumulation of callose in response to *G. orontii* attack (arrows) in non-penetrated (B,C) and penetrated (D,E) cells (open arrows point to the forming haustorium) in Col-0 and *mlo2 mlo6 mlo12* sprayed with 20 µM *β*-estradiol 3 hours before inoculation. Scale bars, 10 μm. All values are mean ±SD. *P* < 0.05, one-way ANOVA and Tukey HSD test, n = 4-5.

The *mlo2 mlo6 mlo12* mutant displayed enhanced callose accumulation following fungal attack, but this response was largely unaffected by inhibition of either RabA or RabE (Fig. 6b,c; Supplementary Fig. 7a,b, Supplementary Video 2). Similarly, robust callose encasements formed around developing haustoria in the successful infections observed (Fig. 6d,e; Supplementary Fig. 7c,d). Thus, RabE-dependent trafficking contributes to both nonhost and *mlo*-mediated powdery mildew immunity independently of major effects on callose deposition. To find support for our observations, we crossed the temperature sensitive *scd1* allele into the *mlo2 mlo6 mlo12* triple mutant. However, we repeatedly failed to find successful penetrations in the *mlo2 mlo6 mlo12* triple mutant and only rarely when in combination with the *scd1* mutation in the restrictive temperature (Supplementary Fig.7e,f). The complete absence of penetrations in the *mlo2 mlo6 mlo12* mutant suggests that an increase in temperature negatively affects the powdery mildew fungus and thereby masks the effect of *scd1*.

## Discussion

Pre-invasive immunity relies on the rapid delivery of defence components to sites of attempted pathogen ingress, yet the trafficking pathways underlying these responses remain poorly defined (Nielsen, 2024). Here, we show that members of the RabA and RabE subfamilies make distinct contributions to plant pre-invasive immunity. RabE-mediated trafficking is required for immunity to powdery mildew fungi, whereas RabA-dependent trafficking contributes predominantly to immunity against the oomycete pathogen *P. infestans*. These findings suggest that pre-invasive immunity is mediated by specialized trafficking pathways rather than a single general defence route.

Several observations support a direct role for RabE-mediated trafficking in powdery mildew immunity. Inhibition of RabE rapidly increased penetration by *B. hordei*, whereas equivalent inhibition of RabA had little effect. Likewise, loss of the RabE GEF SCD1 compromised immunity to *B. hordei*, and RabE inhibition partially overcame the durable immunity conferred by the *mlo2 mlo6 mlo12* triple mutant. Notably, these effects occurred without major reductions in callose accumulation and thus underlines our previous observations that callose accumulation correlates poorly with immunity (Munch et al., 2026). Instead, susceptibility correlated with reduced accumulation of PEN1 at papillae, suggesting that focal delivery of defence-associated membrane components is a key determinant of powdery mildew immunity. The selective involvement of RabE is particularly intriguing given the established connection between RabE proteins and myosin XI-mediated transport (Orr et al., 2021). Previous studies have shown that RabE interacts with myosin XI and functions in polarized secretion, while class XI myosins are required for PEN1 accumulation and pre-invasive immunity to powdery mildew fungi (Yang et al., 2014). The requirement of the barley myosin Required for mlo resistance1 (ROR1) for *mlo*-mediated immunity further strengthens this connection (Acevedo-Garcia et al., 2022). Moreover, RabE interacts with the exocyst complex that mediates secretion but also plays an important role in papilla formation (Pecenková et al., 2011; Ortmannová et al., 2022; Pang et al., 2022). Together, these observations support a model in which RabE, myosin XI and exocyst operate within a common trafficking pathway that promotes localized delivery of defence cargo during powdery mildew attack.

In contrast, the contribution of RabA to powdery mildew immunity appears largely indirect. Dominant-negative RabA2a rapidly altered the localization of multiple plasma membrane proteins, yet defects in pre-invasive immunity to powdery mildew only became apparent following prolonged induction. RabA therefore appears to function primarily in plasma membrane protein homeostasis, with the eventual defects in this form of immunity arising from progressive disruption of cell surface trafficking rather than from inhibition of a dedicated defence pathway. By contrast, short-term inhibition of RabA, but not RabE, significantly increased penetration by P. infestans, indicating that RabA may play a more direct role in pre-invasive immunity against oomycete attack.

Our findings fit well with our recent proposal that pre-invasive immunity comprises at least two genetically separable trafficking pathways (Munch et al., 2026). The present work identifies RabE-and RabA-mediated trafficking as key components of these pathways and links their activity to immunity against distinct groups of filamentous pathogens. We propose that RabE acts together with myosin XI and the exocyst to promote focal delivery of defence cargo during powdery mildew attack, whereas RabA predominantly contributes to plasma membrane protein homeostasis while supporting immunity to oomycete invasion. More broadly, these results suggest that pre-invasive immunity does not rely on a single dedicated defence route but instead emerges from the coordinated action of specialized trafficking modules with distinct cargo specificities and pathogen requirements.

---

**Supplementary Fig. 1:**
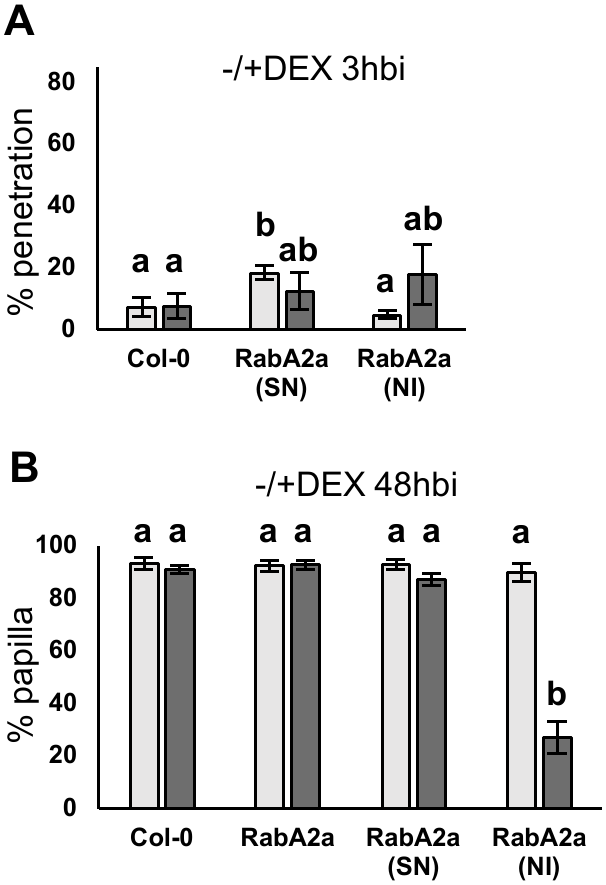
Frequency of successful penetrations of leaf epidermal cells by *B. hordei* in Col-0, *DEX::RabA2a (SN)* or *DEX::RabA2a (NI)* sprayed with either control (white bars) or 20 µM dexamethasone (grey bars) 3 hours before inoculation (A). Frequency of papillae (B) in response to *B. hordei* as detected by staining for callose in in Col-0, *DEX::RabA2a*, *DEX::RabA2a (SN)* or *DEX::RabA2a (NI)* treated as in A. All values are mean ±SD. *P* < 0.05, one-way ANOVA and Tukey HSD test, n = 4-5.

**Supplementary Fig. 2:**
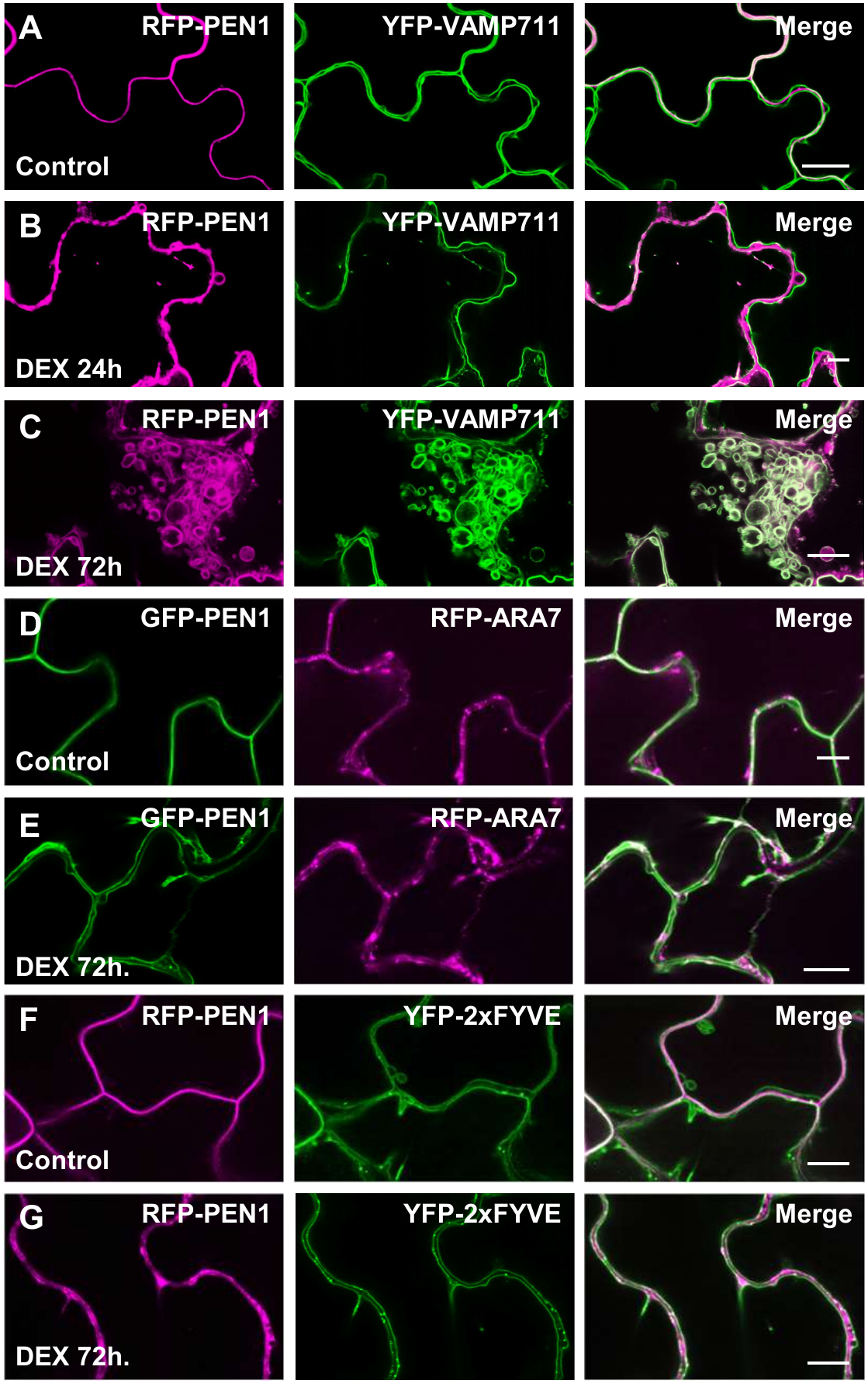
Representative confocal micrographs of epidermal leaf cells from 7-day-old *DEX::RabA2a (NI)* seedlings grown on medium containing 20 μM dexamethasone for the indicated periods before imaging. Localization of RFP-PEN1 and the tonoplast marker YFP-VAM711 following induction of RabA2a (NI) (A-C). Localization of GFP-PEN1 and the MVB marker RFP-ARA7 (D,E). Localization of RFP-PEN1 and the MVB marker YFP-2xFYVE (F,G). Scale bars, 10 μm.

**Supplementary Fig. 3:**
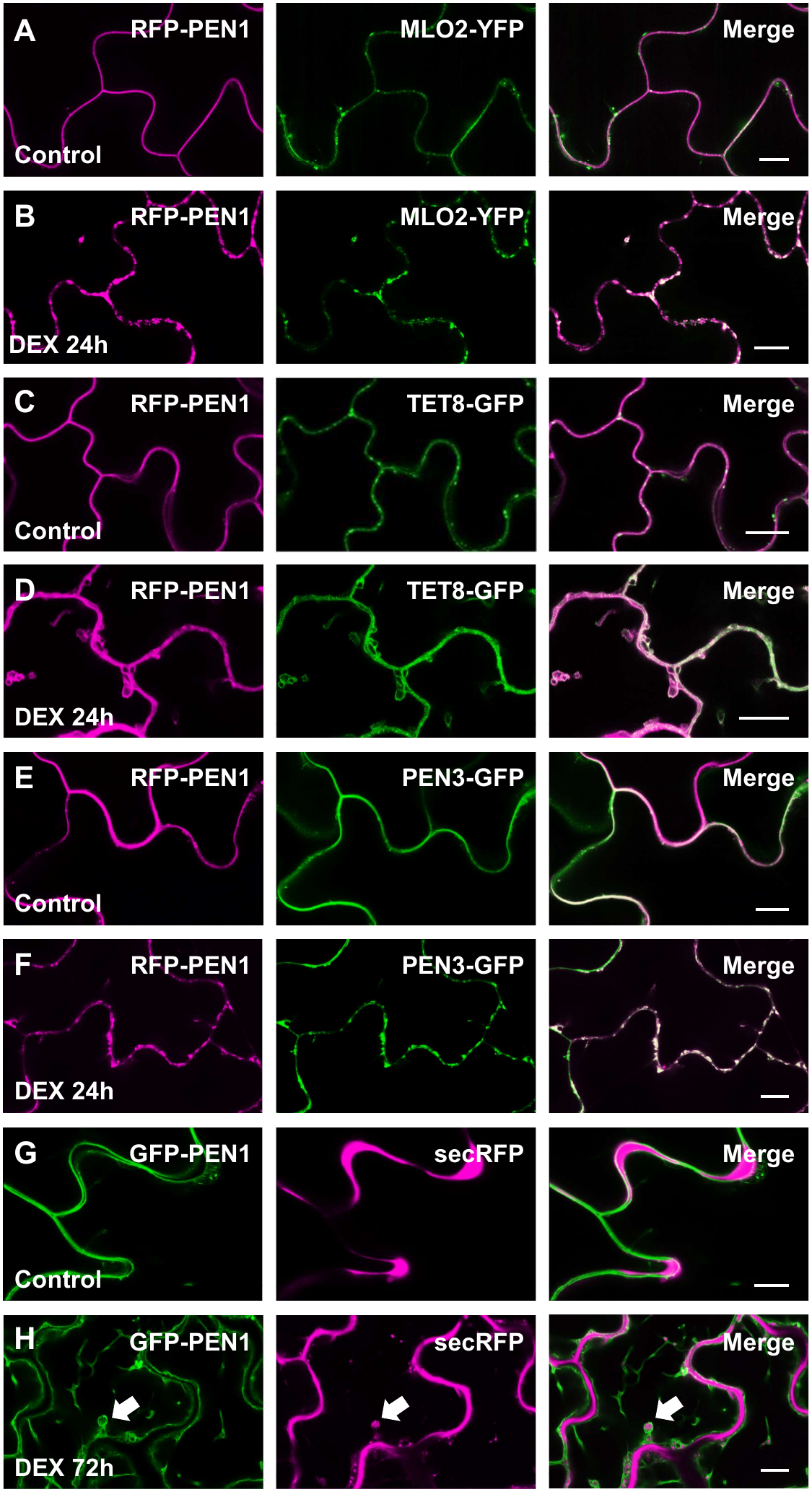
Representative confocal micrographs of epidermal leaf cells from 7-day-old *DEX::RabA2a (NI)* seedlings grown on medium containing 20 μM dexamethasone for the indicated periods before imaging. Localization of RFP-PEN1 and MLO2-YFP (A,B), RFP-PEN1 and TET8-GFP (C,D), RFP-PEN1 and PEN3-GFP (E,F) and GFP-PEN1 and secretory marker secRFP (G,H) following induction of RabA2a (NI). Scale bars, 10 μm.

**Supplementary Fig. 4:**
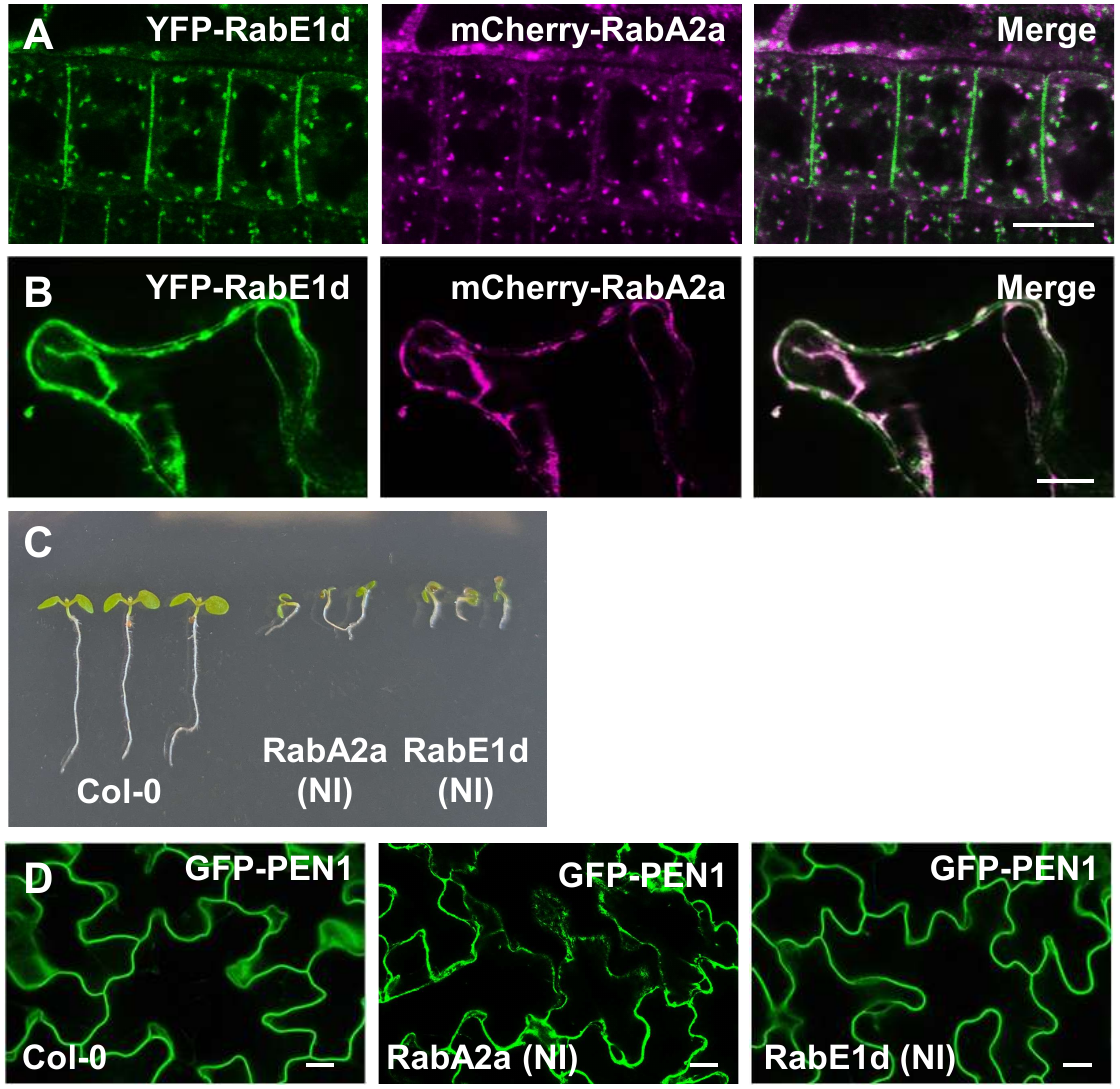
Representative confocal micrographs of root (A) and epidermal leaf cells (B) from plants expressing YFP-RabE1d and mCherry-RabA2a. Scale bars, 10 μm. 7-day-old seedlings of Col-0, *EST::RabA2a (NI)* and *EST::RabE1d (NI)* grown on media supplemented with 10 µM *β*-estradiol (C). Localization of GFP-PEN1 in Col-0, *EST::RabA2a (NI)* or *EST::RabE1d (NI)* sprayed with 20 µM *β*-estradiol 24 hours before imaging (D). Scale bars, 20 μm.

**Supplementary Fig. 5:**
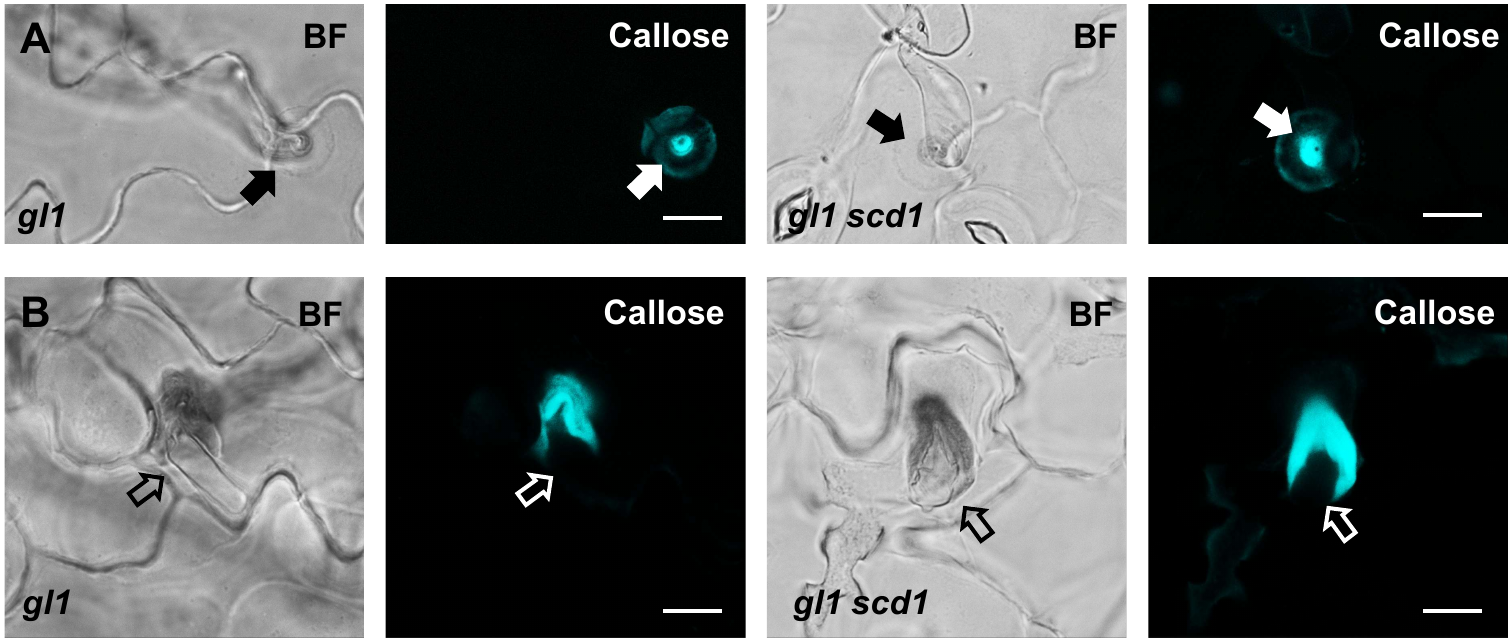
Accumulation of callose in response to *B. hordei* attack (arrows) in non-penetrated (A) and penetrated (B) cells (open arrows point to the forming haustorium) in *glabrous1* (*gl1*) or *gl1 scd1* at 25°C. Scale bars, 10 μm.

**Supplementary Fig. 6:**
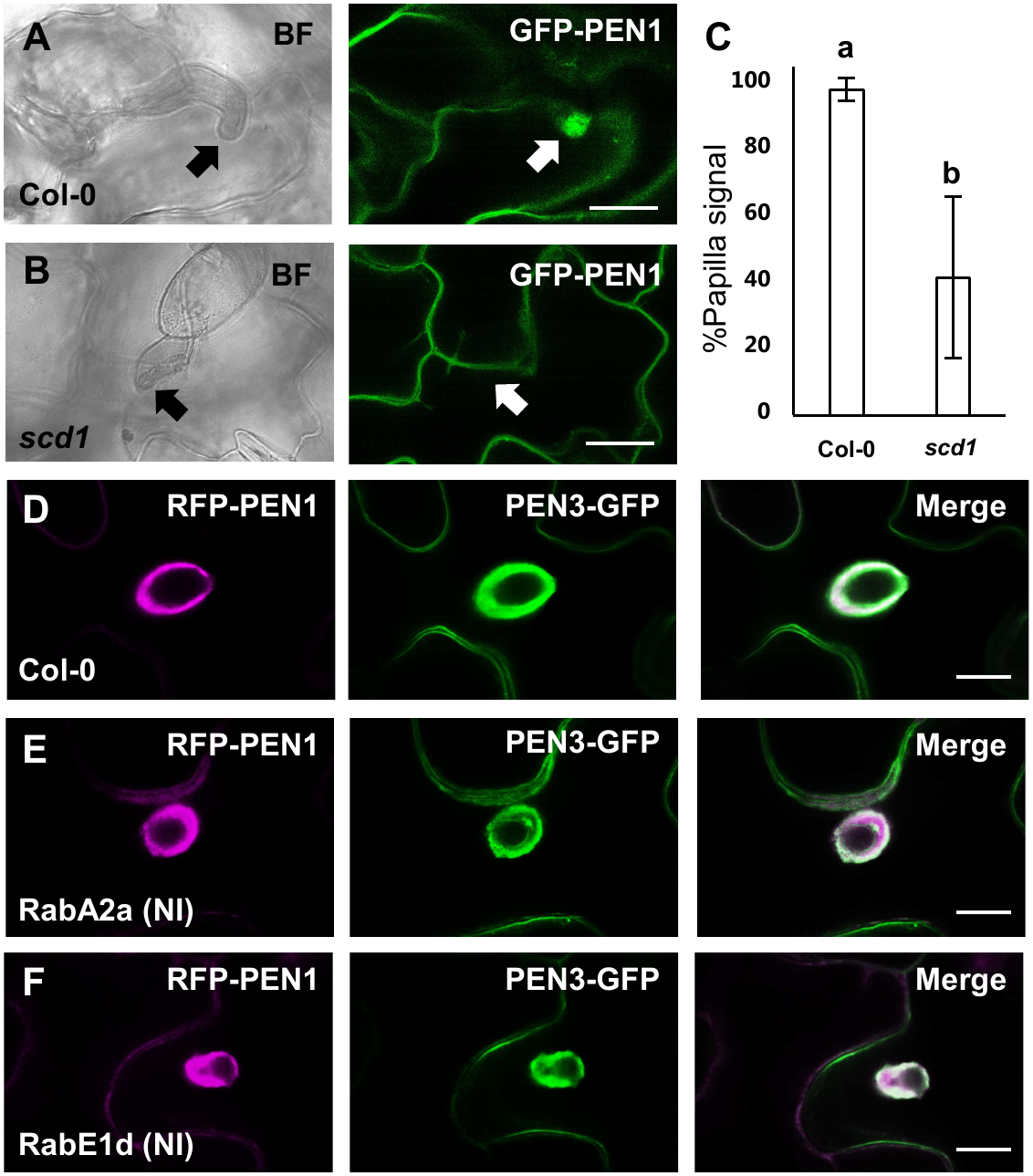
Papillary accumulation of GFP-PEN1 in response to attack by *B. hordei* (arrows) in Col-0 or *scd1* at 25°C (A,B). Frequency of GFP-PEN1 accumulating at *B. hordei* attack sites in Col-0 and *scd1* at at 25°C (C). Accumulation of RFP-PEN1 and PEN3-GFP in encasements in response to successful penetration by *B. hordei* in Col-0, *EST::RabA2a (NI)* or *EST::RabE1d (NI)* sprayed with 20 µM *β*-estradiol 24 hours before imaging (D-E). Scale bars, 10 μm. All values are mean ±SD. *P* < 0.05, one-way ANOVA and Tukey HSD test, n > 86.

**Supplementary Fig. 7:**
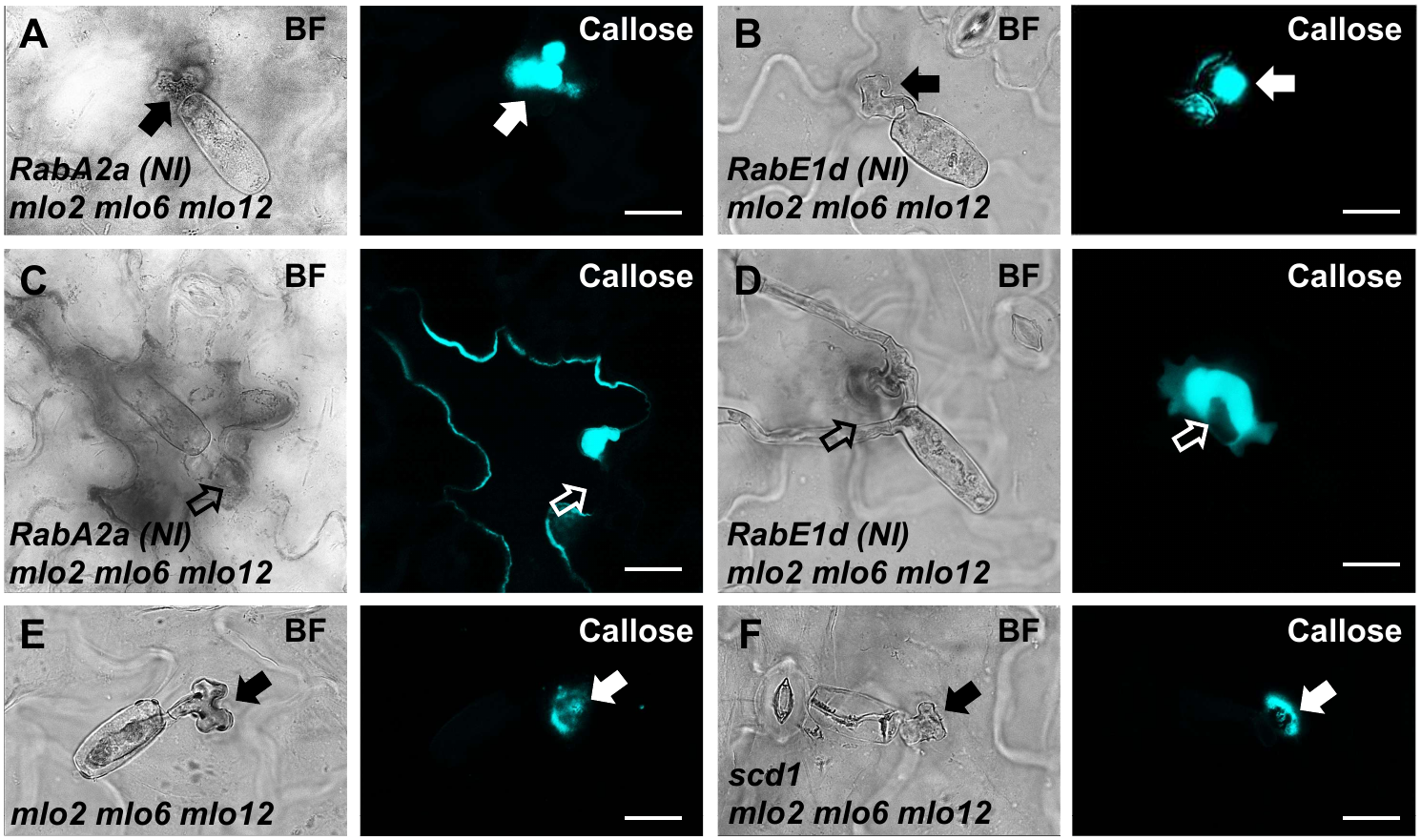
Accumulation of callose in response to *G. orontii* attack (arrows) in non-penetrated (A,B) and penetrated (C,D) cells (open arrows point to the forming haustorium) in *mlo2 mlo6 mlo12* with *EST::RabA2a (NI)* or *EST::RabE1d (NI)* sprayed with 20 µM *β*-estradiol 3 hours before inoculation. In *mlo2 mlo6 mo12* expressing RabA2a (NI) we did not find spores with a fully developed haustorium and secondary hyphal growth. Instead, all penetrated cells had undergone a cell death reaction. In (E,F), accumulation of callose in response to *G. orontii*attack (arrows) in non-penetrated cells in *mlo2 mlo6 mlo12* or *mlo2 mlo6 mlo12 scd1* at 25°C. Scale bars, 10 μm.

**Supplementary Video 1:**

Accumulation of callose in response to *P. infestans* attack in a penetrated cell expressing RabA2a (NI).

**Supplementary Video 2:**

Accumulation of callose in response to *G. orontii* attack in a penetrated cell in *mlo2 mlo6 mlo12*.

## Contributions

HMR, DHM, JM and MEN designed the research, performed experimental work, and analyzed the data. The original draft of the manuscript was written by MEN, with writing, review and editing contributions from all other authors.

## Acknowledgements

The authors would like to thank Mette Sylvan for excellent technical assistance. For generously providing plasmids and seed of mutants and stable transgenic lines we would like to thank Liam Elliot, Gerd Jürgens and Sebastian Y. Bednarek. Imaging data was collected at the Center for Advanced Bioimaging at PLEN. This research was supported by the Novo Nordisk Foundation (NNF19OC0056457 and NNF0086341).

## Materials and Methods

### Plant and fungal growth

Plants of *Arabidopsis thaliana* were grown at 19°C, with 8 h of light at 125 micro Einstein s^−1^ m^−2^ per day. Plants were grown on plates containing 0.5× MS salts (Murashige and Skoog, Duchefa), with 0.8% (w/v) agar (Type M, Sigma) and moved to soil. For transient induction assays, dexamethasone or *β*-estradiol was either supplemented in plates (10 µM) or sprayed (20 µM) onto rosette leaves of 4-week-old plants.

All mutant plant lines were reported previously (*pen1-1*, Collins et al., 2003; *gl1* and *gl1 scd1-1*, Falbel et al., 2003; *mlo2-5 mlo6-2 mlo12-1* and *mlo2-6 mlo6-4 mlo12-8*, Consonni et al., 2006;). Transgenic lines expressing dexamethasone inducible RABA2a WT, N125I, S26N or Q71L (Chow et al., 2008) were obtained from the Eurasian Arabidopsis Stock Centre (uNASC). The following transgenic lines were described previously: GFP-PEN1, RFP-PEN1 and GFP-SYP122 (Rubiato et al., 2022); GFP-SYP132 (Enami et al., 2009); YFP-VAMP711 and YFP-RabE1d (Geldner et al., 2009); RFP-ARA7 (Dettmer et al., 2006); YFP-2xFYVE (Simon et al., 2014); MLO2-YFP (Munch et al., 2026); TET8-GFP (Boavida et al., 2013); PEN3-GFP (Stein et al., 2006); secRFP (Samalova et al., 2006); mCherry-RabA2a (Kirchelle et al., 2016).

The barley powdery mildew fungus, *Blumeria hordei* (isolate C15), and the *A. thaliana* powdery mildew fungus, *Golovinomyces orontii*, were propagated on barley and *A. thaliana*, respectively. *Phytophthora infestans* strain 88069 (A1 mating type, race 1.3.4.7) was maintained on rye agar plates supplemented with 2% sucrose, 100 mg/L ampicillin, 10 mg/L rifampicin.

### Generation of transgenic lines expressing estradiol inducible Rab GTPases

The open reading frame of RabA2a (N125I) was amplified from the DEX::RabA2a (N125) stable line (Chow et al., 2008) and cloned into a Gateway ready pENTR/TOPO-D vector using Multisite Gateway Cloning Technology (Invitrogen). RabE1d (N128I) was generated by a two-step PCR reaction from cDNA and cloned into pENTR/TOPO-D vector. RabA2a (N125I) and RabE1d (N128I) were cloned into a modified *β*-estradiol inducible pMDC7 vector in which Ubiquitin 10 promoter replaced the original promoter (Karnahl et al., 2017). Transgenes *pMDC7::RabA2a (N125)* and *pMDC7::RabE1d (128I)* were transformed into GFP-PEN1, RFP-PEN1 or GFP-PEN1 *mlo2-6 mlo6-4 mlo12-8* and selected for hygromycin resistance (20mg/L) using the floral-dip method (Clough & Bent, 1998).

### Fluorescent protein and callose detection using confocal microscopy

Samples were examined using a ×63 water immersion lens mounted on a Leica Stellaris 8 confocal microscope. For detection and localization of the fluorophores and stains, green fluorescent protein (GFP) was excited at 488 nm and detected at 500–520 nm. Yellow fluorescent protein (YFP) was excited at 508 nm and detected at 515–575 nm. mCherry was excited at 590 nm and detected at 600–650 nm. Aniline blue-stained callose was excited at 440 nm and detected at 450–500 nm. Imaging data were collected at the Center for Advanced Bioimaging, University of Copenhagen (Denmark). Projections of serial confocal sections, image overlays, and contrast enhancement were performed using the image processing software (GIMP 2).

### Infection studies

Spores from *B. hordei* and *G. orontii* were blown evenly onto the leaves of 4-to 5-week-old *A. thaliana* plants and kept at 19°C. Zoospores of *P. infestans* were released from sporangia by addition of cold water and incubation at 4°C for 90 min. The zoospore suspension was diluted to 5 × 10^5^ or 1 × 10^6^ spores/mL and added to the abaxial side of attached leaves from 4-week-old *A. thaliana* plants. The leaves were kept in a closed Petri dish on water-soaked filter paper at 19°C. For infections studies on *scd1* mutants, plants were moved to 25°C immediately after inoculation.

For scoring penetration success, leaf material was Trypan blue (T6146; Sigma-Aldrich) stained at 24 h post inoculation for *B. hordei*, 48 h post inoculation for *G. orontii*, and 24 h post inoculation for *P. infestans*. Penetration success was determined by light microscopy based on the presence of a developing haustorium inside epidermal cells. Once the penetration status for a particular event was determined, only then was the callose response investigated. Callose staining was performed using 0.01% Aniline blue (415 049; Sigma-Aldrich) in 1 M glycine (adjusted to pH 9.5 using NaOH) and visualized by UV epifluorescence. Statistics are based on > 50 attack sites per leaf of four to five individual plants.

### Primers

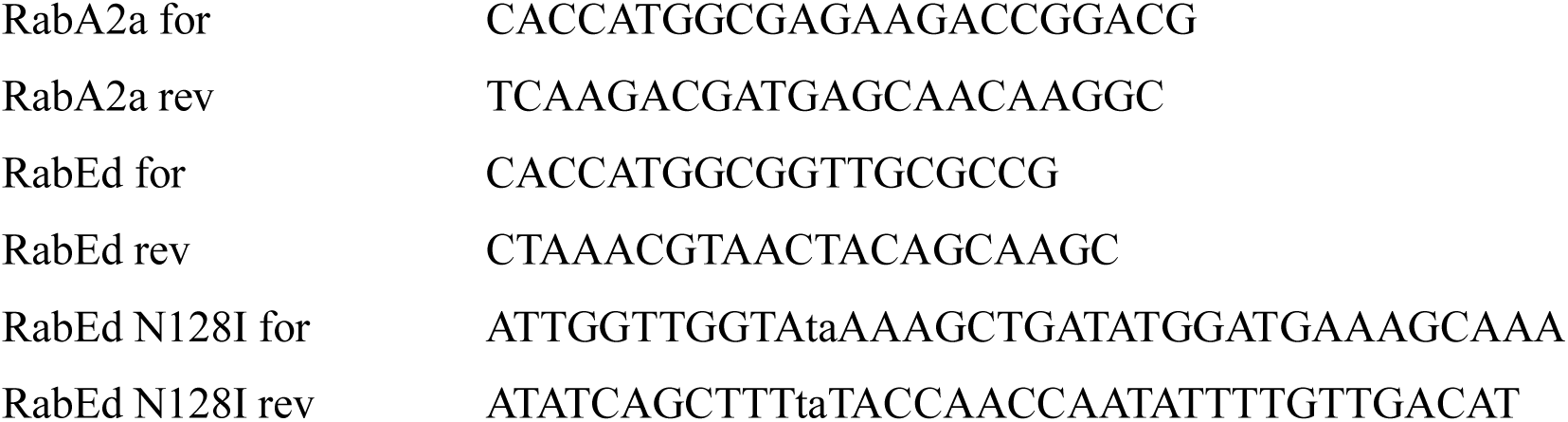

